# Decoding the interconnected splicing patterns of hepatitis B virus and host using large language and deep learning models

**DOI:** 10.1101/2025.07.28.667110

**Authors:** Chun Shen Lim, Chris M. Brown

## Abstract

Hepatitis B virus (HBV) infection causes approximately one million deaths annually and remains a major driver of hepatocellular carcinoma. Despite its compact 3.2-kb genome, HBV exhibits extensive alternative splicing. Functionally, HBV splice variants contribute to immune evasion and reduce the likelihood of achieving a functional cure. Here, we show that HBV splicing efficiency—quantified from 279 RNA-seq libraries of HBV-associated liver biopsies and cultured cells—correlates more strongly with disease progression than the overall proportion of spliced HBV RNAs, an emerging biomarker. All HBV splice sites are embedded within protein-coding regions, forming a genetic architecture distinct from typical host splice sites. To decode the sequence determinants of HBV splicing, we apply transformer-based and deep learning models to 4,707 HBV genomes. These models reveal that more highly used HBV splice sites are more conserved and share features with host splice sites that are less frequently used but still functional. This similarity likely reflects constraints imposed by HBV’s compact genome, which must accommodate overlapping protein-coding regions. HBV may have evolved to exploit suboptimal but spliceable host-like motifs without disrupting its genetic architecture. Further analysis of splicing propensity across HBV genomes reveals genotype-specific patterns, indicating regulation by sequence context in a site- and genotype-dependent manner. HBV genotypes may have coevolved with their human hosts to fine-tune splicing through host-like features, supporting mechanisms of viral persistence and immune evasion. This study demonstrates the utility of AI in decoding viral splicing architectures and provides a framework for investigating co-transcriptional processes in other clinically important viruses.

**Impact Statement:** Hepatitis B virus (HBV) is a major global health concern, causing 1.1 million deaths in 2022 and 1.2 million new infections each year. It is a leading cause of serious liver conditions, including cirrhosis and liver cancer. Although vaccines can prevent HBV infection, there is currently no cure. HBV produces different types of genetic messages (RNAs), including spliced versions that are processed by the host cell’s machinery. These spliced RNAs help the virus evade the immune system and make it harder for treatments to fully clear the infection. In this study, we analysed 279 HBV samples from liver tissues and lab-grown cells and found that the efficiency of host-mediated splicing of viral RNA reflects the severity of disease. Using advanced artificial intelligence tools, we mapped the splicing patterns in both the virus and human host, and investigated over 4,700 HBV genomes. We discovered that HBV splice sites resemble host splice sites that are used less frequently but remain functional, suggesting the virus has evolved RNA sequences that are compatible with the host cell’s splicing machinery while accommodating its compact genome. Insights into this viral adaptation may help researchers identify new biomarkers for disease severity and develop therapeutic strategies that disrupt the virus’s ability to exploit the host cell’s machinery.

**Data Summary:** The raw RNA-sequencing (RNA-seq) libraries analysed in this study were previously published and are available in the Gene Expression Omnibus under accession number GSE155983. These libraries form part of a curated collection of 279 RNA-seq datasets derived from HBV-associated liver biopsy tissues and cultured cells. Further details and metadata are available in the associated GitHub repository: https://github.com/lcscs12345/HBV_splicing_paper_2025.

## INTRODUCTION

Chronic hepatitis B virus (HBV) infection remains a major global health challenge and the leading cause of hepatocellular carcinoma (HCC)^1^. Current antiviral therapies suppress viral replication rather than eliminate the virus, allowing it to persist in infected hepatocytes. HBV continues to drive nearly half of all HCC cases worldwide, with the Asia-Pacific region bearing a disproportionate burden, accounting for over 70% of HBV-related HCC deaths^2–4^. With liver cancer incidence projected to exceed one million cases annually by 2025^1^, there is an urgent need to better understand the viral sequence determinants that contribute to disease progression and treatment resistance, to inform precision strategies for monitoring and intervention.

HBV is a DNA retro-transcribing virus with a compact, 3.2-kb genome. HBV produces over 20 spliced RNA variants through the use of splice sites embedded within the protein-coding regions of its pregenomic RNA (pgRNA)^5–7^. Alternative usage of these splice sites can give rise to a variety of functional HBV protein isoforms. For example, the hepatitis B spliced protein (HBSP), derived from the SP1 splice variant, promotes immune escape by reducing immune cell recruitment and suppress apoptosis via PI3K/Akt activation^8,9^. In contrast, hepatitis B doubly spliced protein (HBDSP), derived from SP7, functions as a transactivator that promotes apoptosis in HCC cells^10,11^. The polymerase-surface fusion protein (P-S FP), derived from SP13, is a structural protein that may substitute for the large HBV surface protein^12^. The viral oncogenic transactivator HBx, encoded by an intact X open reading frame (ORF) retained in most HBV splice variants, plays a key role in transcriptional regulation and hepatocarcinogenesis^5^.

Notably, HBV splice variants can be packaged into defective viral particles and detected in patient serum^13–15^. Although these particles are incapable of autonomous replication, elevated levels of HBV splicing in serum correlate with progression of chronic liver disease^14,15^. Such splicing activity can be observed years before HCC diagnosis, even in patients with normal alpha-fetoprotein tumour marker levels^16^. These observations support the potential HBV splice variants as biomarkers of disease progression.

HBV is classified into at least ten genotypes (A–J), each with distinct geographic distributions and sequence characteristics that affect clinical outcomes^17–19^. These genotypes differ in their association with disease progression, treatment response, and risk of HCC. We previously demonstrated that the major HBV genotypes express markedly distinct compositions of splice variants^6^, suggesting that regulation of splicing may be genotype-specific. HBV splicing can be regulated both in *cis*, for example by the post-transcriptional regulatory element (PRE)^20^, and in *trans*, through host factors such as DDX5 and DDX7^21^. A recent study showed that DDX5 and DDX7 RNA helicases can suppress recognition of an HBV splice donor site^21^. However, the sequence determinants and genotype-specific features that control splice site selection and usage in HBV remain poorly understood, particularly in the context of disease progression and genotype-specific variation.

In this study, we employ large language and deep learning models (SpliceBERT and OpenSpliceAI)^22,23^ to systematically analyse HBV splice site usage across 279 curated viral transcriptomes^6^ and 4,707 viral genomes from HBVdb^19^. To better interpret splicing patterns, we introduce two terms—’CDS splice sites’ and ‘typical splice sites’. CDS splice sites are located within protein-coding regions, while typical splice sites occur at the boundaries between protein-coding exons and introns. This distinction allows direct comparisons between HBV and host splicing architecture. Our findings reveal that HBV splicing is shaped by sequence context, host-like features, and viral genotype, offering insights into potential implications for viral replication, immune evasion, oncogenesis, and biomarker development for precision strategies.

## RESULTS

### Splicing patterns in HBV transcriptomes reflect disease state

We previously assembled 279 HBV transcriptomes by analysing 513 publicly available RNA-seq libraries derived from HBV-associated liver biopsy tissues and cultured cells^6^. These transcriptomes revealed highly complex splicing patterns across the major HBV genotypes^6^ (Supplementary Fig S1). Notably, all HBV splice sites identified are splice sites located within protein-coding exons, which we term ‘CDS splice sites’.

To assess the clinical relevance of HBV splicing, we compared the proportions of spliced HBV RNAs across different sample types. The median proportion of spliced HBV RNAs ranged from 3% to 22%. Advanced-stage HCC samples, as characterised by portal vein tumour thrombosis (PVTT), exhibited significantly higher levels of HBV splicing than earlier-stage tumours and adjacent liver tissues (Fig 1A). HBV-infected primary human hepatocytes (PHHs) and HBV-transfected Huh7 hepatoma cells also showed elevated splicing levels, likely influenced by experimental conditions, such as the use of a strong CMV promoter for expressing HBV pregenomic transcripts (pgRNAs)^6^.

**Fig 1.**
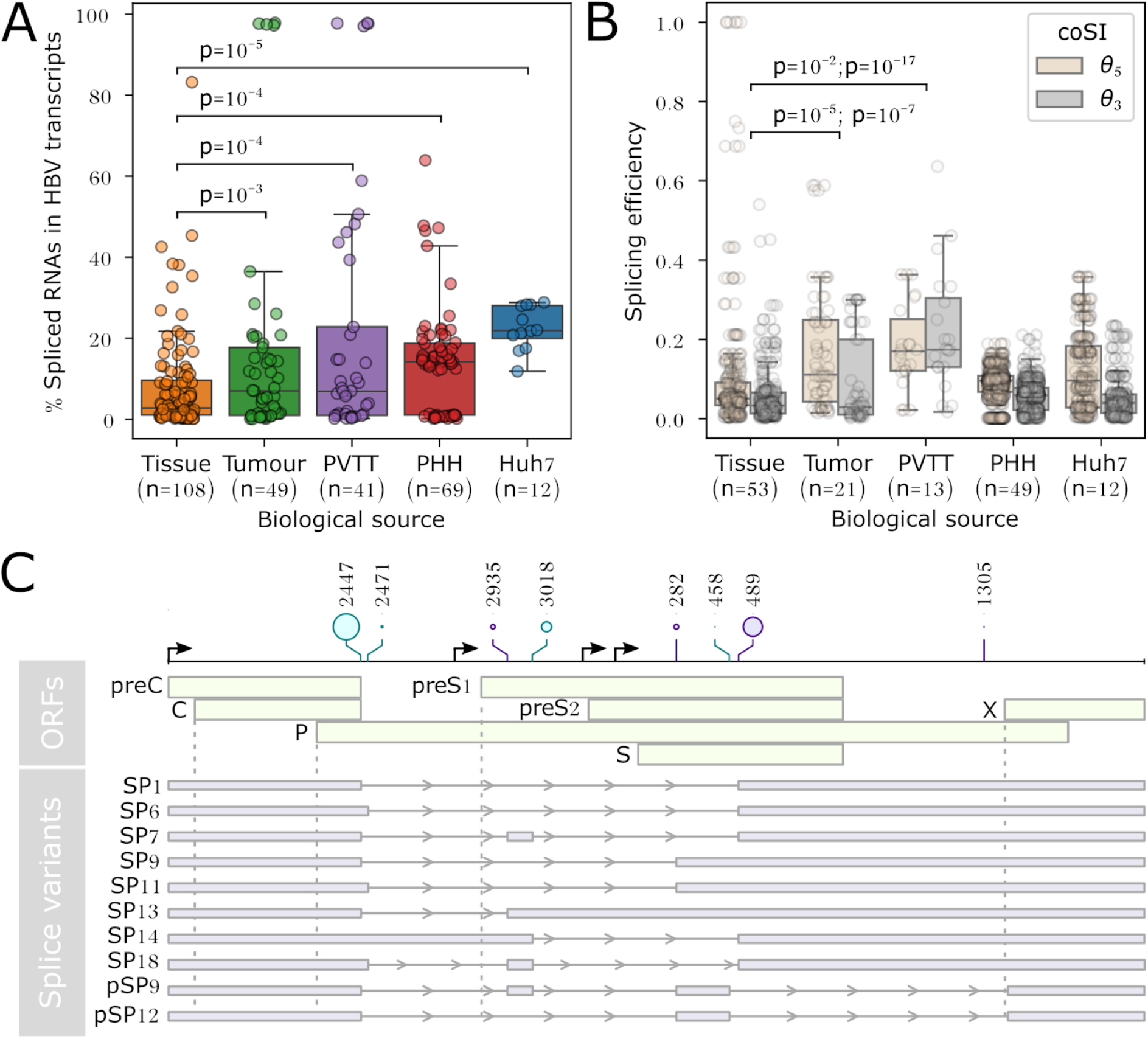
The proportion of spliced HBV RNAs varies across liver tissue types and disease progression. **(A)** Liver tissues adjacent to tumours show the lowest proportions of spliced HBV RNAs, followed by tumours, advanced-stage PVTT (portal vein tumour thrombosis), HBV-infected primary human hepatocytes (PHHs), and HBV-transfected Huh7 cells. P-values are derived from independent t-tests. **(B)** Across splice variants, donor sites generally exhibit higher splicing efficiency than acceptor sites, as indicated by higher completed splicing index (coSI) θ_5_ scores compared to θ_3_ scores. **(C)** The top 10 most common splice variants identified across 279 HBV transcriptomes. All splice sites are located within the protein-coding regions of the virus open reading frames (ORFs). Lollipop sizes are scaled by the median coSI, ranging from 0.0060 to 0.21.

Further analysis of HBV transcriptomes revealed that samples from tumours and PVTT exhibited significantly higher splicing efficiency than PHHs and Huh7 cells, as measured by the completed splicing index (coSI)^24,25^ (Fig 1B). This suggests that splice site-level measurement may serve as a more specific biomarker of disease progression than overall spliced RNA proportions, which has been proposed as an emerging biomarker^14–16^. Although HBV splice sites are generally weak, the most commonly expressed splice variants derive from donor sites with higher splicing efficiency than acceptor sites, with the exception of SP6 (Fig 1C, Supplementary Fig S1A). Genotype C was the most prevalent genotype in the dataset and showed substantially greater splicing activity than genotype B (Supplementary Fig S1B).

### HBV splice sites are distinct from typical host splice sites

As multiple lines of evidence suggest that HBV leverages the host splicing machinery during cancer progression, we sought to decipher and compare the splicing code between the virus and host. To decode the viral and host splicing architectures, we harnessed the representational power of SpliceBERT, the state-of-the-art transformer-based large language model (LLM)^23^. We fine-tuned SpliceBERT using the Spliceator training set^26^ to capture conserved splicing patterns across 114 species, a strategy shown to outperform models trained solely on *Homo sapiens* data^23^. We then compared HBV splice sites (44 donor and 98 acceptor sites from four representative transcriptomes) with host splice sites (95,323 donor and 99,179 acceptor sites) by extracting nucleotide embeddings and applying PCA (Principal Components Analysis), UMAP (Uniform Manifold Approximation and Projection), and Leiden hierarchical clustering. Host splice site data were derived from plus-strand transcripts to streamline this analysis, which should be considered when interpreting downstream analyses such as gene ontology enrichment.

While host splice sites and non-splice sites (non-donor GU and non-acceptor AG motifs) formed distinct clusters, HBV splice sites were distributed across multiple clusters and exhibited 4–6 times lower normalised mutual information (NMI) scores than host splice sites, indicating greater heterogeneity (Fig 2). Furthermore, many HBV splice sites were grouped into clusters containing a mix of splice and non-splice sites—a “grey zone” in the embedding space (Fig 2B, D). This unexpected pattern led us to hypothesise that HBV splice sites in these mixed clusters may be less efficiently recognised by the splicing machinery, as indicated by their cryptic sequence contexts.

**Fig 2.**
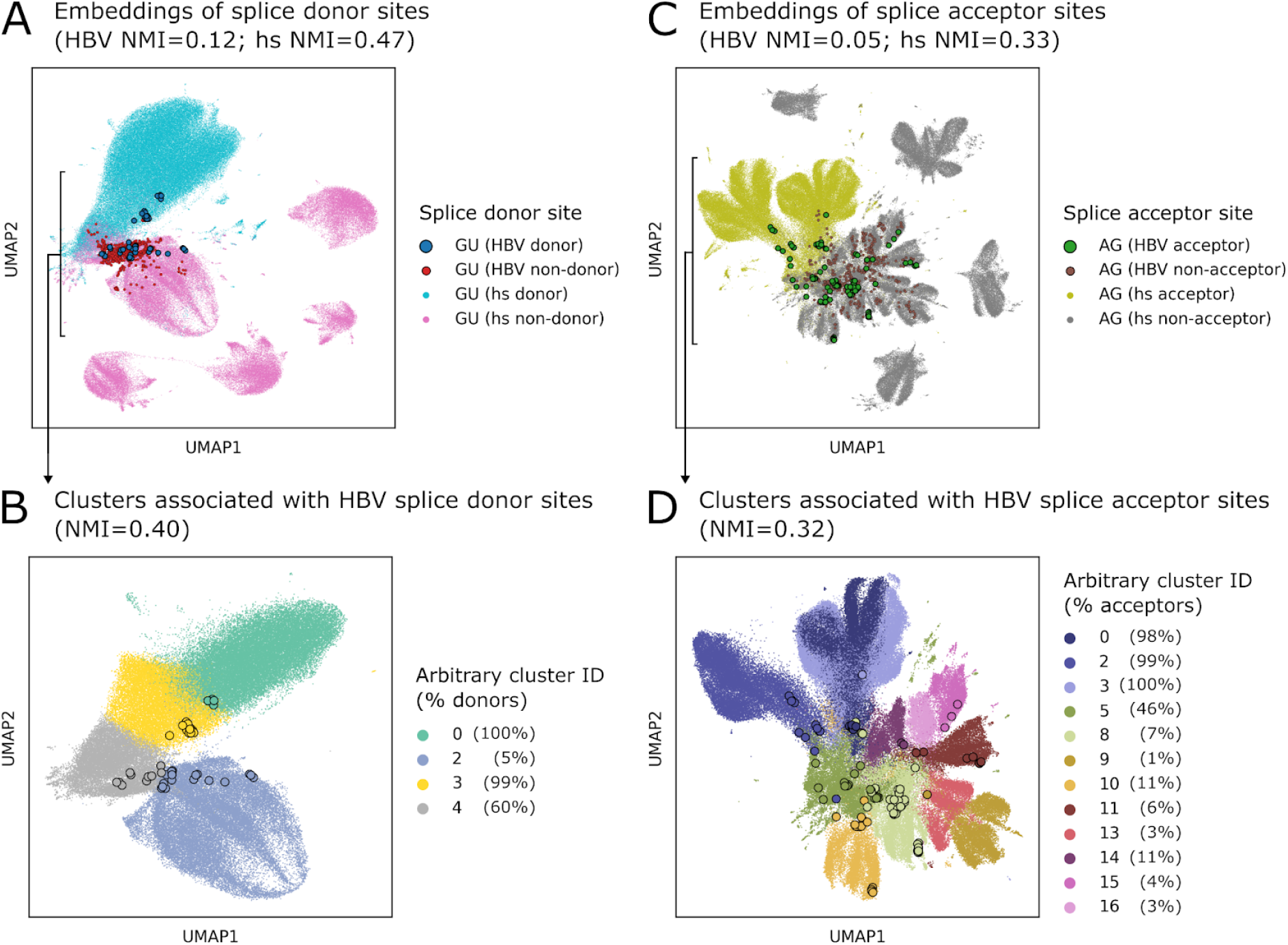
Most HBV splice sites are distinct from host splice sites. Clustering of nucleotide embeddings from HBV and *Homo sapiens* (hs) splice donor (**A** and **B**) and acceptor (**C** and **D**) sites, extracted from the final layer of SpliceBERT’s splice donor model. Points for HBV splice sites are outlined in black. Embeddings were subjected to dimensionality reduction using PCA followed by UMAP, and clustered using the Leiden algorithm. Normalised mutual information (NMI) scores indicate that HBV splice sites are more distinct from host splice sites. This analysis includes 44 HBV donor and 98 acceptor sites, along with 624 non-donor GU sites and 525 non-acceptor AG sites, derived from four randomly selected HBV transcriptomes among HBV associated samples. Host splice sites include 95,323 donor and 99,179 acceptor sites, along with 80,542 non-donor GU sites and 123,162 non-acceptor AG sites.

To test this hypothesis, we mapped experimentally determined splicing efficiency scores (coSI) from HBV-expressing Huh7 cells^6^ to the Leiden clusters. We found that splicing efficiency positively correlates with the proportion of true, annotated splice sites within each cluster (Fig 3A, Supplementary Fig S2; Spearman’s correlation 0.8, p-value 10^−4^). This supports our idea that splice sites which are more efficiently recognised and processed by the splicing machinery tend to cluster together, while weakly recognised sites, such as HBV splice sites, are more likely to co-cluster with non-splice sites.

**Fig 3.**
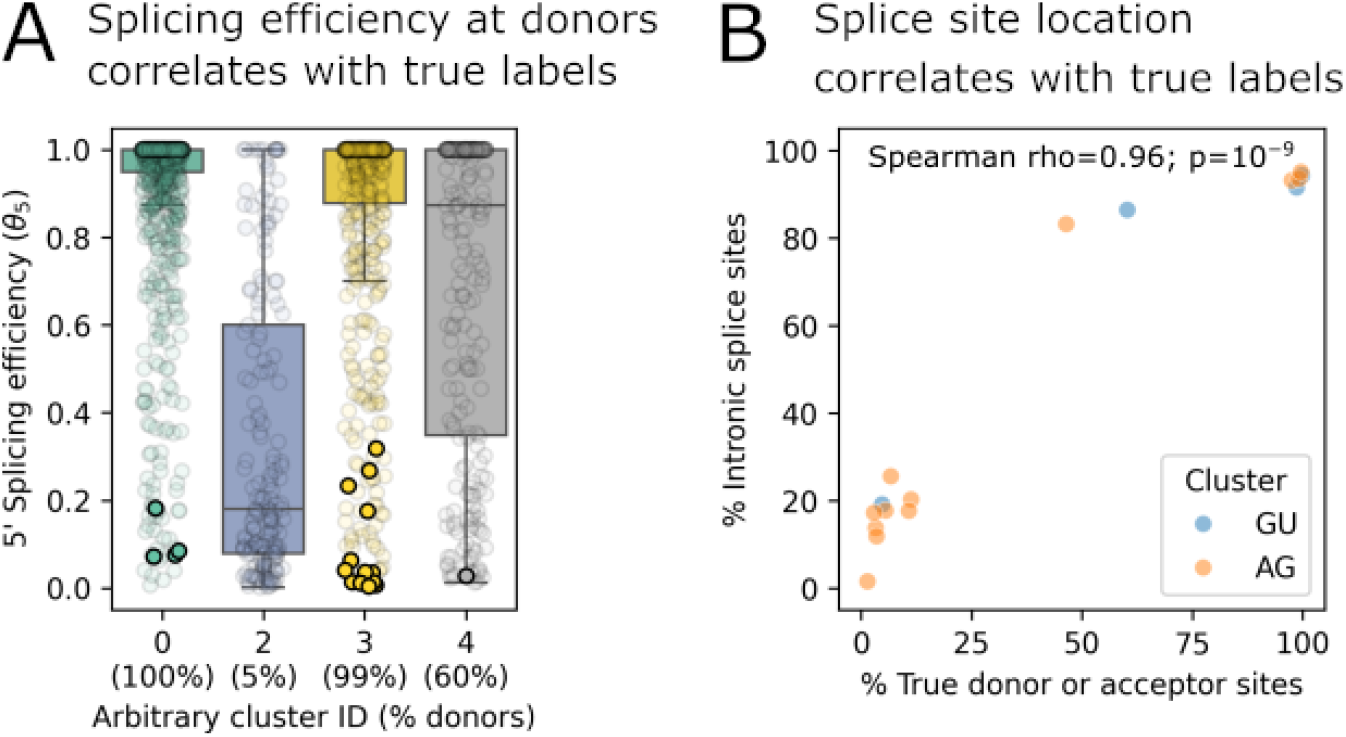
Splicing efficiency (A) and splice site location (B) correlate with true label proportions in Leiden clusters. Points for HBV splice sites are outlined in black (**A**). Leiden clusters were derived from the embedding space of splice sites shown in Fig 2, where HBV and host splice sites were grouped based on nucleotide embeddings from SpliceBERT. Percent typical splice sites was calculated from the proportion of splice sites located exclusively at the boundaries between protein-coding exons and introns. Clusters enriched for true, annotated splice sites tend to show higher splicing efficiency and a greater proportion of typical splice sites, supporting the biological relevance of the embedding-based clustering. See Supplementary Fig S2 for the splicing efficiency at the acceptor sites and true label proportions in Leiden clusters.

We further examined the distribution of splice sites located exclusively at the boundaries between protein-coding exons and introns (typical splice sites) across these clusters. Consistent with the above results, clusters with higher proportions of true, annotated splice sites also contained more typical splice sites (Fig 3B). In contrast, CDS splice sites, which are embedded within protein-coding regions, are more likely to fall into mixed clusters and have a lower splicing efficiency. This supports the idea that typical splice sites are more efficiently processed, while CDS splice sites are more likely to evade recognition due to their cryptic sequence contexts.

These findings suggest that HBV’s compact genome imposes structural constraints, requiring it to accommodate overlapping coding regions and essential RNA elements. To maintain functionality under these constraints, HBV appears to exploit host-like motifs that are suboptimal yet spliceable, enabling splicing without disrupting its genetic architecture.

Notably, a similar configuration is observed in a small subset of host genes that contain both CDS donor and acceptor splice sites. These host genes are rare, comprising only 3% and 1% of all CDS splice donor and acceptor sites, respectively. This subset is enriched for the Gene Ontology term “organelle organization” (GO:0006996; adjusted p-value 10^−7^), although it should be noted that this analysis was restricted to plus-strand transcripts.

### AI models reveal genotype- and site-specific splicing propensities

Our analyses suggest that splice site recognition by the host splicing machinery is highly context-dependent. However, it remains unclear whether these splicing patterns are conserved across HBV genotypes or shaped by genotype-specific sequence features. Our previous transcriptomic study supports the latter^6^, suggesting that splicing propensity may be influenced by genotype-dependent sequence variation.

To further investigate genotype-specific splicing propensities, we performed a sliding window analysis across 4,707 HBV genomes, including 1,728 genotype B, 2,084 genotype C, 252 genotype F, and 643 recombinant forms (RF)^19^. Each 400-nucleotide window was scored using SpliceBERT. Sequence-level representations were extracted from the classification token [CLS] of SpliceBERT, which captures global contextual information for each input sequence. These sequence-level representations were subjected to dimensionality reduction (PCA and UMAP) and clustering using the Leiden algorithm. This analysis revealed distinct clusters corresponding to HBV genotypes, indicating that splicing propensity is influenced by genotype-specific sequence features (Fig 4).

**Fig 4.**
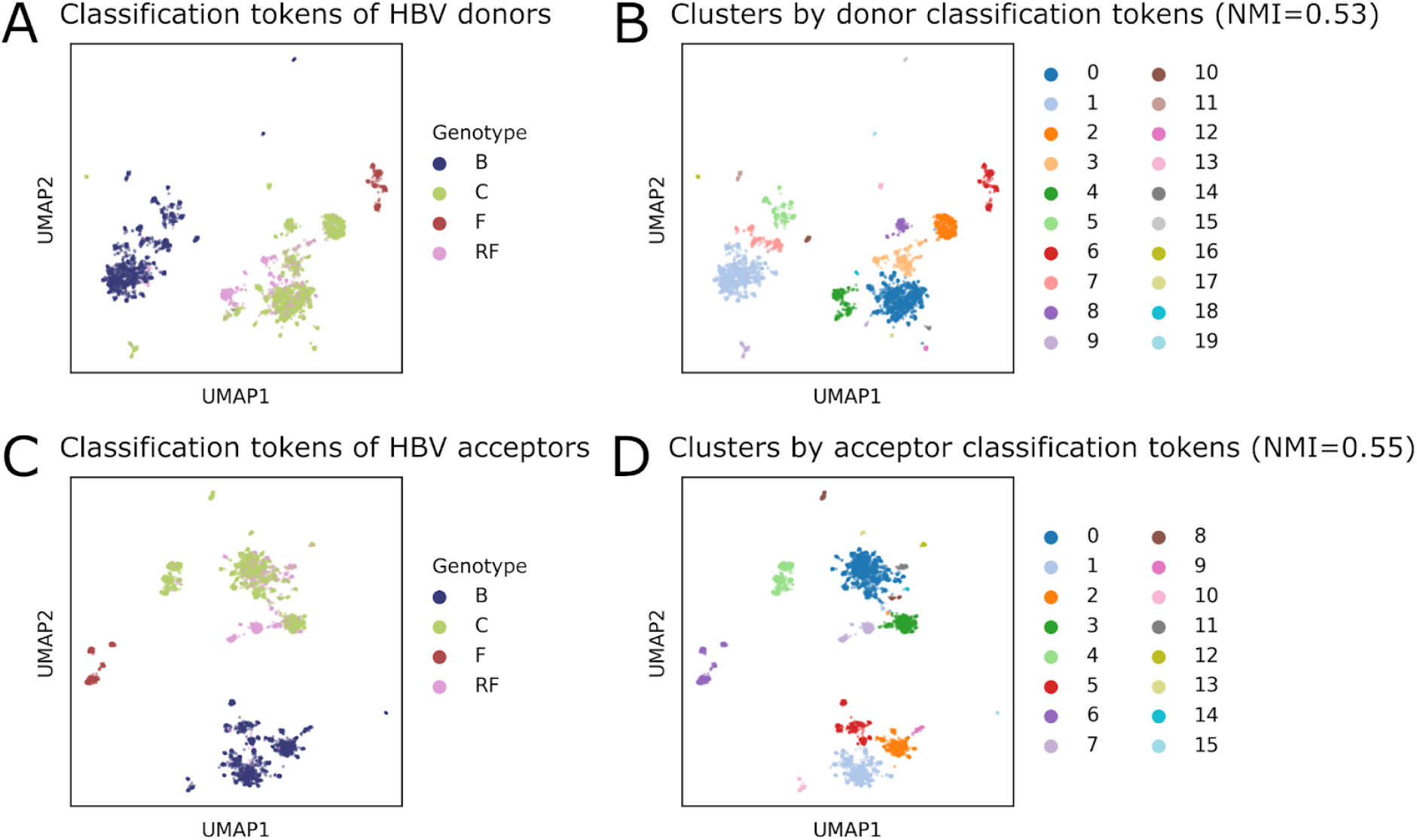
Splicing propensities are distinct between HBV genotypes. A sliding window of 400 nucleotides with 1-nt overlap was applied across 4,707 HBV genomes, including 1,728 genotype B, 2,084 genotype C, 252 genotype F, and 642 recombinant forms (RF). Each window was scored using SpliceBERT, and the resulting [CLS] token outputs were subjected to dimensionality reduction using PCA followed by UMAP, and clustered using the Leiden algorithm.

We next examined whether HBV splice sites possess sequence features that distinguish them from non-splice sites, and whether these features correlate with splice site usage. To address this, we utilised SpliceBERT and OpenSpliceAI, the latter being a deep convolutional neural network trained using *H. sapiens* data^22^. Although these models were trained for binary classification, we grouped HBV splice sites into higher and lower usage categories based on coSI scores. Higher usage splice sites consistently scored higher than lower usage and non-splice sites (Fig 5), indicating that frequently used sites possess more recognisable sequence features.

**Fig 5.**
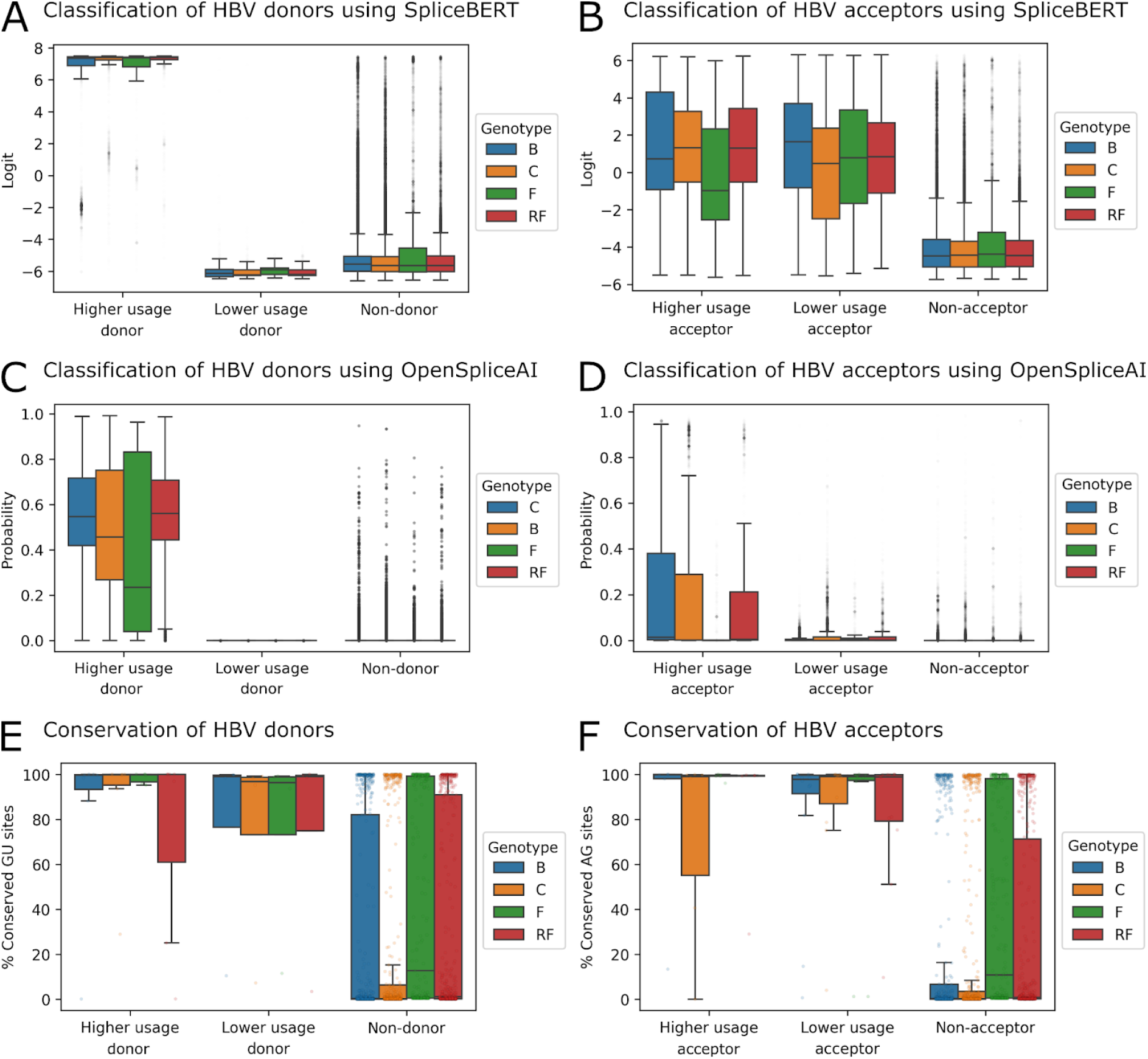
HBV splice donor and acceptor sites are conserved and distinct from non-splice sites. Splice sites were categorised by usage level based on coSI scores (≥0.1 for higher usage, <0.1 for lower usage). Higher usage splice sites exhibit higher SpliceBERT [CLS] scores (logit values; panels **A** and **B**) and OpenSpliceAI prediction probabilities (panels **C** and **D**), along with stronger splice site motif conservation compared to lower usage and non-splice sites (panels **E** and **F**). In contrast, lower usage splice sites generally show low scores similar to non-splice sites, except for SpliceBERT scores at lower usage acceptor sites (panel **B**). Sites were selected from a diverse set of HBV genomes, comprising 1,728 genotype B, 2,084 genotype C, 252 genotype F, and 642 RF. These included seven higher usage and four lower usage donor sites, six higher usage and 12 lower usage acceptor sites, along with 1,011 non-donor GU sites and 943 non-acceptor AG sites. Refer to Fig 4 for splicing propensity analysis using SpliceBERT [CLS] scores.

However, distinguishing lower usage splice sites, particularly donor sites, proved more challenging (Fig 5A and C). This difficulty was reflected in both models, in conjunction with notable differences in scoring across HBV genotypes. Prediction scores for donor splice sites varied substantially between genotypes, suggesting that genotype-specific sequence variation affects how well these sites are scored by AI models trained on *H. sapiens* or multi-species splicing data. SpliceBERT performed better for acceptor site prediction, while OpenSpliceAI showed stronger performance for donor sites (Table 1), highlighting the complementary strengths of these models.

**Table 1.** Performance of AI models in predicting HBV splice sites. Metrics include area under the ROC curve (AUROC), area under the precision-recall (AUPRC), F1 score, precision, and recall, evaluated using curated HBV genomic datasets. See related Figs 4 and 5.

| AI Model |  | AUROC | AUPRC | F1 score | Precision | Recall |
| --- | --- | --- | --- | --- | --- | --- |
| SpliceBERT | Donor | 0.72 | 0.59 | 0.51 | 0.45 | **0.59** |
|  | Acceptor | **0.90** | **0.60** | **0.63** | 0.63 | **0.62** |
| OpenSpliceAI | Donor | **0.78** | **0.60** | 0.51 | **0.99** | 0.34 |
|  | Acceptor | 0.89 | 0.57 | 0.13 | **0.97** | 0.07 |

Note: The best results are marked in bold. This analysis includes 40,117 donor sites, 757,682 non-donor GU sites, 67,262 acceptor sites, and 652,160 non-acceptor AG sites, derived from 2,084 genotype C, 1,728 genotype B, 252 genotype F, and 642 recombinant forms (RF).

Although these models were not trained on viral data, the higher scores assigned to more frequently used HBV splice sites suggest that AI models can identify functional sequence features resembling host splice sites that are less frequently used. This capability enables us to probe the HBV genetic architectures and uncover genotype-specific regulatory patterns. For example, the elevated SpliceBERT scores for higher usage splice sites in genotype C align with the more extensive splicing observed in liver issues infected with this genotype (Fig 5A, B, and Supplementary Fig S1B). While the development of virus-specific models may improve predictive accuracy, the priority remains experimental validation to establish the clinical relevance and therapeutic potential of these predictions.

### Higher usage splice sites are conserved

As HBV splice site usage varies across genotypes and higher usage sites tend to be more distinguishable by AI models, we next asked whether these frequently used splice sites are also more conserved across HBV genomes. If so, this would suggest that higher usage splice sites are functionally important and subject to selective pressure, distinguishing them from lower usage and non-splice sites.

To test this, we computed the proportion of canonical intron motifs (GU for donor sites and AG for acceptor sites) across 4,707 HBV genomes. In general, higher usage splice sites were more conserved, whereas lower usage splice sites were less conservation, and non-splice sites exhibited highly variable motif frequencies (Fig 5E and F). Notable exceptions included higher usage donor sites in RF and higher usage acceptor sites in genotype C, which were less conserved than their counterparts in other genotypes.

To further investigate the sequence features that may influence splice site recognition, we analysed base frequencies around HBV donor and acceptor sites grouped by usage level. For comparison, we included *H. sapiens* splice sites located within protein-coding regions (CDS splice sites) and those located exclusively at the boundaries between protein-coding exons and introns (typical splice sites). Overall, HBV splice sites with higher usage shared more similar sequence contexts with host CDS splice sites than with lower usage HBV splice sites (Fig 6). For example, HBV acceptor sites with higher usage show polypyrimidine tracts more similar to those observed in host CDS acceptor sites than host typical acceptor sites (Fig 6 and Supplementary Fig S3). Our sequence conservation analysis suggests that sequence context determines splice site selection and usage, and that HBV may exploit host-like sequence features to control splicing. These parallels suggest that HBV has evolved to mimic functional, though infrequently used, host splicing architectures to facilitate immune evasion and persistence.

**Fig 6.**
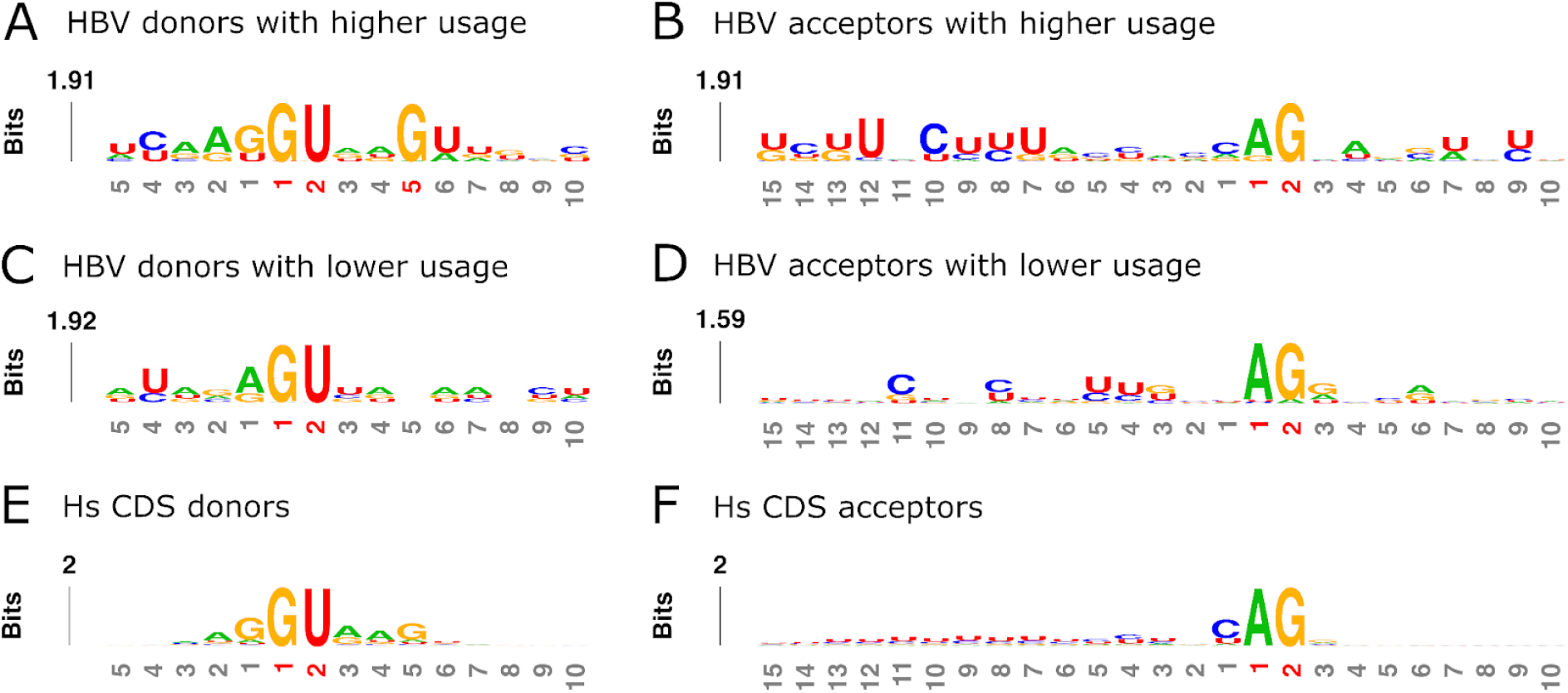
HBV donor and acceptor splice sites with higher usage exhibit sequence contexts similar to those splice sites found within *H. sapiens* protein-coding regions. Sequence logos illustrate HBV donor sites with higher and lower usage (panels **A** and **C**; both 18,828 sites), HBV acceptor sites with higher and lower usage (panels **B** and **D**; 23,535 and 51,777 sites, respectively), *H. sapiens* donor and acceptor sites (panels **E** and **F**; 1,243 and 646 sites, respectively) located within the protein-coding regions of 20,852 mRNAs. The consensus polypyrimidine tracts observed at the HBV acceptor sites with higher usage (panel **B**) are more similar to those observed in host CDS acceptor sites (panel **F**) than to host typical acceptor sites (see Supplementary Fig S3B).

## DISCUSSION

Our findings demonstrate that HBV splicing is not transcriptional noise but a regulated process with functional consequences in disease progression. We showed that HBV splicing levels are elevated in liver tumours and advanced-stage PVTT samples (Fig 1), consistent with a role in hepatocarcinogenesis. Previous studies have shown that these splice variants can be packaged into defective virus particles^13–15^. These defective particles are detectable in patient serum, and their overall proportion has been proposed as an emerging biomarker^14,15^. Notably, the splice site-level indicator shown in this study may act as a more specific and mechanistically informative biomarker. These HBV splice variants and their encoded proteins have also been shown to control viral replication dynamics^27^, suppress host innate immune response, such as IFN-α signaling^28^, and are associated with a reduced chance of achieving functional cure^14^. A functional cure is a clinical endpoint where there is a sustained loss of hepatitis B surface antigen (HBsAg) and HBV DNA after a finite course of treatment^29^.

To dissect the sequence-function relationship of HBV splice sites, we leveraged 279 HBV transcriptomes derived from deep RNA-seq datasets^6^, 4,707 HBV genomes curated from HBVdb^19^, and state-of-the-art AI models, including SpliceBERT large language model^23^ and OpenSpliceAI deep learning framework^22^. This integrative approach enabled us to identify splice sites with high usage and predictive sequence features. We found that higher usage splice sites are more readily recognised by AI models, exhibit greater conservation across HBV genomes, and share sequence characteristics with host CDS splice sites, despite the constraint imposed by the virus’s compact genome. These converging lines of evidence suggest that HBV may co-opt host-like sequence motifs to regulate splicing genotypically, thereby enhancing its persistence and immune evasion^7^.

Collectively, our study provides a comprehensive framework for understanding HBV splicing through the lens of over 50,000 years of virus-host coevolution, highlighting a sophisticated level of host mimicry^17,30,31^. By applying cutting-edge computational tools to large-scale genomic and transcriptomic data, we highlight the conserved, mechanistically important, and clinically relevant nature of HBV splicing. These findings lay the groundwork for precision therapeutic targeting of HBV splice variants and their regulatory elements, offering new avenues for intervention in chronic HBV infection and HBV-related HCC, including the development of novel CRISPR-based therapies^32^. Furthermore, our findings demonstrate the broader utility of AI models in decoding viral genomic architectures and regulatory mechanisms, with implications for studying splicing and post-transcriptional control in other clinically important viruses.

## METHODS

### Data

We curated a total of 279 HBV transcriptomes from our previous transcriptomic study^6^. The dataset includes BAM and SJ.out.tab files generated by STAR v2.7.6a in 2-pass mode^33^, and Gene Transfer Format (GTF) annotation files produced by StringTie v1.3.3b^34^. For *H. sapiens*, UCSC Genome Browser’s hg19 genome and GENCODE annotation v41lift37 were used^35,36^.

### Splicing efficiency analysis

Splicing efficiency scores for donor and acceptor sites (coSI θ_5_ and θ_3_ scores, respectively) were computed from BAM files using IPSA^25^. To map splice variant IDs to their respective coSI scores, GTF files were converted to BED file format using UCSC Genome Browser’s KentUtils, and intersected with IPSA output using BEDTools v2.31.1^37^.

### Embedding-based clustering

To capture conserved splicing patterns across eukaryotic species, we fine-tuned SpliceBERT using the Spliceator training set, generating model weights for donor and acceptor sites^23,26^. We modified SpliceBERT fetch_embedding.py to incorporate fine-tuned weights into the pretrained 510-nt model using HuggingFace Transformers v4.27.2^38^. Input files were prepared by generating HBV pgRNA consensus sequences from BAM files using BCFtools v1.22^39^. Splice junctions supported by ≥5 uniquely mapping reads were extracted from SJ.out.tab files and converted to BED format.

For *H. sapiens*, splice sites located on the plus strand were extracted to streamline the analysis. Non-donor GU sites and non-acceptor AG sites within protein-coding regions were used as negative controls. Viral and host data were combined, and the final layer of nucleotide embeddings was extracted for all splice sites, representing local sequence context. All model inference and embedding extraction were performed on NVIDIA A100 40G GPUs.

### Splicing propensity analysis

To compare splicing propensity across HBV genotypes, we applied a sliding window of 400-nt with 1-nt overlap for 4,707 HBV genomes from HBVdb^19^. All genomes chosen for this analysis were of the same length to prevent misalignment at splice sites. Genomic sequences were rearranged to match the structure of pgRNA, and flanking regions of 200-nt upstream of the preC start codon and downstream of the X stop codon were appended to provide biological context and minimise artificial sequence padding. Each window was scored using the fine-tuned SpliceBERT models. Dimensionality reduction was performed on the sequence-level representations extracted from the [CLS] token using PCA and UMAP, followed by clustering with the Leiden algorithm implemented in SCANPY v1.10.3^40^.

### Splice site classification

We applied SpliceBERT and OpenSpliceAI v0.0.4^22^ to classify HBV splice sites versus non-splice sites. For OpenSpliceAI, we used the 400-nt flanking-size model trained on *H. sapiens* MANE data. For visual comparison, splice sites were grouped into higher usage (coSI>=0.1) and lower usage (coSI<0.1) categories.

### Sequence conservation and logo analysis

To assess splice site motif conservation, we calculated the proportion of conserved GU and AG motifs across HBV genomes. Flanking sequences of splice sites were extracted and visualised using kpLogo v1.1^41^.

### Statistical analysis and data visualisation

All data processing and statistical analyses were performed using pandas v2.2.2^42^ and NumPy v1.26.4^43^. NMI scores were calculated using scikit-learn v1.6.1^44^ to quantify the similarity between clustering results and known splice site labels. Model performance metrics, including area under the receiver operating characteristic curve (AUROC), area under the precision-recall curve (AUPRC), F1 score, precision, and recall, were also computed using scikit-learn. These metrics were used to evaluate the ability of SpliceBERT and OpenSpliceAI to distinguish HBV splice sites from non-splice sites across curated genomic datasets. Independent t-tests and Spearman’s correlation analyses were conducted using SciPy v1.13.1^45^ to assess statistical significance and relationships between the proportion of spliced HBV RNAs, splicing efficiency scores, and clustering patterns. All visualisations were generated using Matplotlib v 3.6.3^46^ and seaborn v0.13.2^47^.

## Abbreviations

AI: artificial intelligence
AUROC: area under the ROC curve
AUPRC: area under the precision-recall
CDS: protein-coding sequence
coSI: completed splicing index
HBDSP: hepatitis B doubly spliced protein
HBSP: hepatitis B spliced protein
HBsAg: hepatitis B surface antigen
HBV: hepatitis B virus
HCC: hepatocellular carcinoma
NMI: normalised mutual information
ORF: open reading frame
PCA: Principal Components Analysis
pgRNA: pregenomic RNA
PHH: primary human hepatocyte
P-S FP: polymerase-surface fusion protein
PRE: post-transcriptional regulatory element
PVTT: portal vein tumour thrombosis
RF: recombinant form
RNA-seq: RNA sequencing
UMAP: Uniform Manifold Approximation and Projection.

## Code and data availability

Scripts and data for the study can be found at https://github.com/lcscs12345/HBV_splicing_paper_2025.

## Author Contributions

C. S. L. conceived the study, acquired funding, conducted the analysis, and wrote the manuscript. C. M. B. acquired funding, reviewed, and edited the manuscript.

