## Supplementary for "Decoding the interconnected splicing patterns of hepatitis B virus and host using large language and deep learning models"

### SUPPLEMENTARY MATERIALS

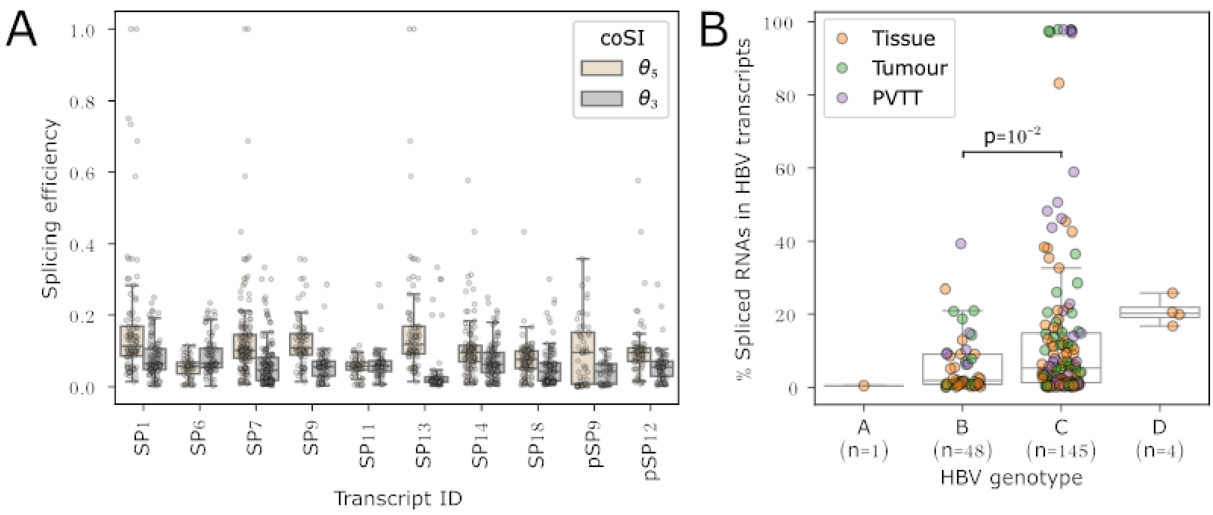


**Fig S1. Splicing efficiency is transcript- and genotype-specific. (A)** HBV splice variants generally exhibit higher splicing efficiency at donor sites than acceptor sites, except for SP6, as indicated by higher completed splicing index (coSI) $\theta_5$ scores compared to $\theta_3$ scores. These transcripts represent the top 10 most common splice variants identified across 279 HBV transcriptomes. **(B)** Samples associated with genotype C show a significantly higher proportion of spliced HBV RNAs compared to those with genotype B.

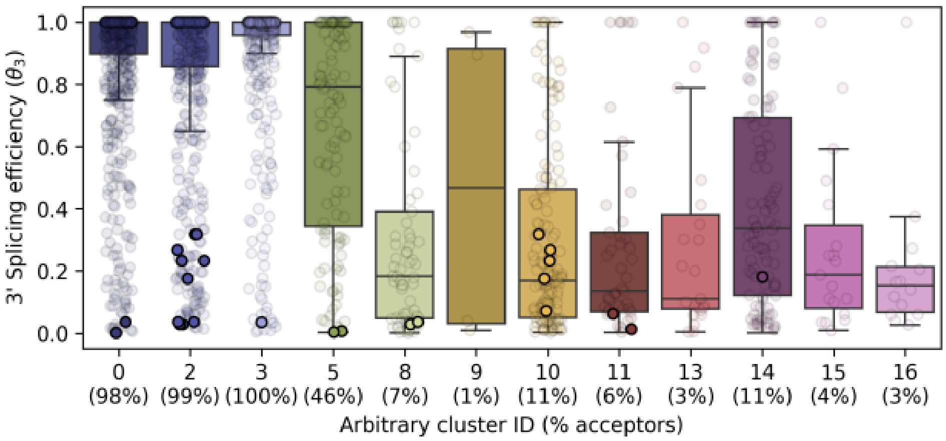


**Fig S2. Splicing efficiency at acceptor sites correlates with true label proportions in Leiden clusters.** Points for HBV splice sites are outlined in black. Related to Figs 2 and 3.

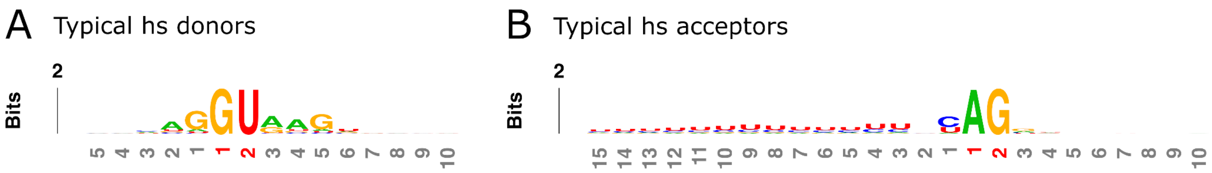


**Fig S3. Sequence contexts of splice sites located exclusively at the boundaries between protein-coding exons and introns in *Homo sapiens*.** Sequence logos represent 37,656 typical splice donor sites **(A)** and 75,312 typical splice acceptor sites **(B)**, derived from 58,963 protein-coding transcripts.
